# Phytoplankton traits from long-term oceanographic time-series

**DOI:** 10.1101/148304

**Authors:** Crispin M Mutshinda, Zoe V Finkel, Claire E Widdicombe, Andrew J Irwin

## Abstract

Trait values are usually extracted from laboratory studies of single phytoplankton species, which presents challenges for understanding the immense diversity of phytoplankton species and the wide range of dynamic ocean environments. Here we use a Bayesian approach and a trait-based model to extract trait values for four functional types and ten diatom species from field data collected at Station L4 in the Western Channel Observatory. We find differences in maximum net growth rate, temperature optimum and sensitivity, half-saturation constants for light and nitrogen, and density-dependent loss terms across the functional types. We find evidence of very high linear loss rates, suggesting that grazing may be even more important than commonly assumed and differences in density-dependent loss rates across functional types, indicating the presence of strong niche differentiation among functional types. Low half-saturation constants for nitrogen at the functional type level may indicate widespread mixotrophy. At the species level, we find a wide range of density-dependent effects, which may be a signal of diversity in grazing susceptibility or biotic interactions. This approach may be a way to obtain more realistic and better-constrained trait-values for functional types to be used in ecosystem modeling.

## Introduction

Phytoplankton perform about half of global photosynthesis, form the base of the marine food web and are important drivers of biogeochemical cycles (Field et al. 1998). Model projections of changes in phytoplankton primary production with climate over the next century are extremely variable (Finkel et al. 2010, Finkel 2014). Projections of changes in communities and biogeochemical cycling usually depend on mechanistic models of phytoplankton productivity parameterized with traits of phytoplankton species (Le Quéré et al. 2005, Litchman et al. 2006). The traits used in models vary according to the research questions, but most commonly include maximum growth rate, Arrhenius-like temperature effects on growth rate, half-saturation parameters linking the growth rate to resource availability, and grazing susceptibility (Litchman et al. 2007, Irwin & Finkel 2016). At present, many of these parameters are not well constrained for phytoplankton communities (Anderson 2005, Irwin & Finkel 2016).

Phytoplankton are evolutionarily and ecologically diverse and include many phyla and tens of thousands of species (Sournia et al. 1991, de Vargas et al. 2015). This complexity presents several challenges for trait-based modeling. Trait values measured in the lab are almost always determined for a few key species, while their applications in models of natural communities usually apply to dozens to thousands of species. The aggregation of similar species into biogeochemically defined functional types greatly simplifies models, but there is no clear way to decide which species should be used as representatives of each functional type (Merico et al. 2004, Le Quéré et al. 2005, Hood et al. 2006). Trait values for species in the same functional type and trait values used in models vary widely, commonly by a factor of 10-100 (Anderson 2005, Irwin & Finkel 2016). It is not clear how to average trait values across species to represent a functional type since phytoplankton growth rate is a non-linear function of trait values. Furthermore, species well adapted to lab conditions may not be representative of their respective functional types growing in natural communities. A second set of challenges concerns the difficulty of using lab-based estimates of trait values in a field context. Trait values quantified using laboratory cultures under controlled conditions are stable under repeated measurement, but there is a challenge in identifying the most appropriate conditions for culture experiments. For example, the maximum growth rate is commonly estimated in the lab, but differences in culture conditions from one lab to another imply that there is always some doubt about the true maximum growth rate for a species (Boyd et al. 2013). Trait-values, including maximum growth rate and nutrient uptake rates, estimated in the field can differ substantially from those measured in the lab (Furnas 1991, Laws 2013, Lomas et al. 2014). Cultures grown under equilibrium conditions in the lab may not reveal key acclimation traits or the consequences of environmental variability that can be crucial to the fate of phytoplankton in natural communities (Grover 1991, Raven 2011). In summary, trait values for most phytoplankton species are not available and we do not currently have enough data to strongly constrain trait values used in functional type models (Anderson 2005, Flynn et al. 2015).

An approach that addresses many of these challenges for determining trait values for functional types is to estimate those values from long-term time series of natural communities observed in the field. Our goal is to obtain quantitative estimates of trait values that define the dynamics of the biomass of phytoplankton functional types. These trait values will be affected by the species that are present in the community, the range of environmental conditions observed, the spectrum of environmental variability, as well as abiotic and biotic interactions. We call them realized traits in recognition that they are not the fixed traits of a particular species. This label is an echo of the difference between fundamental and realized niches, where the realized niche is measured in a community and can differ from the fundamental niche (Hutchinson 1957, Colwell & Rangel 2009). Here we obtain realized trait values by fitting a model of biomass dynamics to time series of phytoplankton functional type biomass and coincident environmental conditions. The model describes temporal biomass changes in terms of net growth rate modified by temperature, irradiance, total available nitrogen concentration, and a density dependent loss term. Realized trait values estimated from field data may be quite different from trait values obtained in the lab and may vary across communities in different locations. The advantage of these realized traits compared to species-level traits quantified in the lab is that these traits by definition describe observed community dynamics.

## Methods

### Data

We used data from the Western Channel Observatory (WCO) oceanographic time-series (www.westernchannelobservatory.org.uk) in the Western English Channel. The WCO data include phytoplankton, zooplankton, and fish trawls together with measurements of several physical and chemical environmental parameters such as temperature, salinity and nutrient concentrations. The data used here were collected at Station L4 (50° 15.00′N, 4° 13.02′W) located about 10 km south of the Plymouth breakwater with a water column depth of about 50 m (Harris 2010). We used 349 weekly observations of taxonomically resolved phytoplankton abundance, temperature, nitrate, nitrite, and ammonium concentrations sampled at 10 m depth in the upper mixed layer and sea-surface irradiance collected over a 7-year period spanning 15 April 2003 through 31 December 2009. Average biovolume measurements were recorded for each species (Widdicombe et al. 2010) and converted to carbon content (Menden-Deuer & Lessard 2000) to obtain biomass concentrations (mg C m^−3^) for each species. We used observations of 193 taxonomic categories identified as 138 species, 27 genera, and 28 size-classes for broader morphological categories. Biomass concentrations were aggregated into four functional types: diatoms, dinoflagellates, coccolithophorids, and phytoflagellates. The phytoflagellate type is taxonomically diverse but is dominated (more than 50% of the biomass) by unidentified flagellates less than 5 μm in diameter. Some species may be benthic or tychoplanktonic. We added together the concentrations of nitrate, nitrite, and ammonium to obtain a single inorganic nitrogen (mg m^−3^) concentration. Most of the variation in total nitrogen concentration is due to variation in nitrate concentration. Irradiance (mol m^−2^ d^−1^) was measured continuously above the sea-surface near Station L4 at Plymouth and averaged over the day. Data for missing weeks were imputed by linear interpolation using the na.approx function from the zoo library in R (R Core Team 2016).

### The model

We describe the multiplicative growth rate of each functional type’s biomass as the product of the following 5 components: (i) a net growth rate reduced by limitation due to either low light or low nitrogen concentration, (ii) a temperature effect, (iii) a density feedback term dependent on the biomass of the focal functional type, (iv) a density feedback term dependent on the biomass of all phytoplankton not in the focal functional type, and (v) a positive multiplicative noise term. The change in biomass from one week to the next (from week *w*–1 to week *w*) for each functional type *i* is modeled by multiplying the biomass in week *w*–1 by the (multiplicative) growth rate according to a stochastic Gompertz model (Saitoh et al. 1997, Mutshinda et al. 2009, Mutshinda et al. 2011). We chose to model the net growth rate as a linear combination of density-independent growth rate and density-dependent losses, which is most appropriate given the lack of direct information about grazing rates, grazer biomass, or viral abundance. Therefore, the biomass concentration *Y_i,w_* (in mg C m^−3^) of the *i*^th^ functional type for each week after the first (*w* ≥ 2) is described by
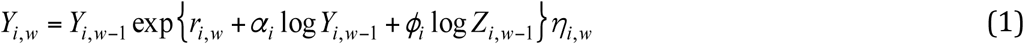

where *Z_i,w_* is the combined biomass concentration of all phytoplankton not including the *i*^th^ functional type during week *w*. The growth rate, which appears in the exponent of Eq. (1), is composed of a density-independent component, *r_i,w_*, and a density-dependent component *α_i_* log(*Y*_*i,w*−1_) + *ϕ_i_* log(*Z*_*i,w*−1_). Stochastic noise enters the biomass dynamical model (Eq. 1) through the random multiplicative noise terms *η*_*i,w*_ > 0 assumed to be serially independent and log-normally distributed with median one and mean 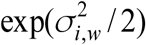, so that the natural logarithms, log(*η_i_*_,*w*_), are independently zero-mean normal with respective variances 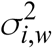. The species-specific error variance 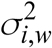 lumps together sampling error and the variability due to other potentially important factors not included in the model. Sampling error can be reduced by using replicated samples. The log-normal distribution adopted here is widely used to describe species abundance and biomass patterns (MacArthur 1960, Sugihara 1980) on both theoretical and empirical grounds. In an earlier study (Mutshinda et al. 2016), we found phytoplankton biomass at this site to be well described by the log-normal distribution. The notation is summarized in Table 1.

**Table 1.**
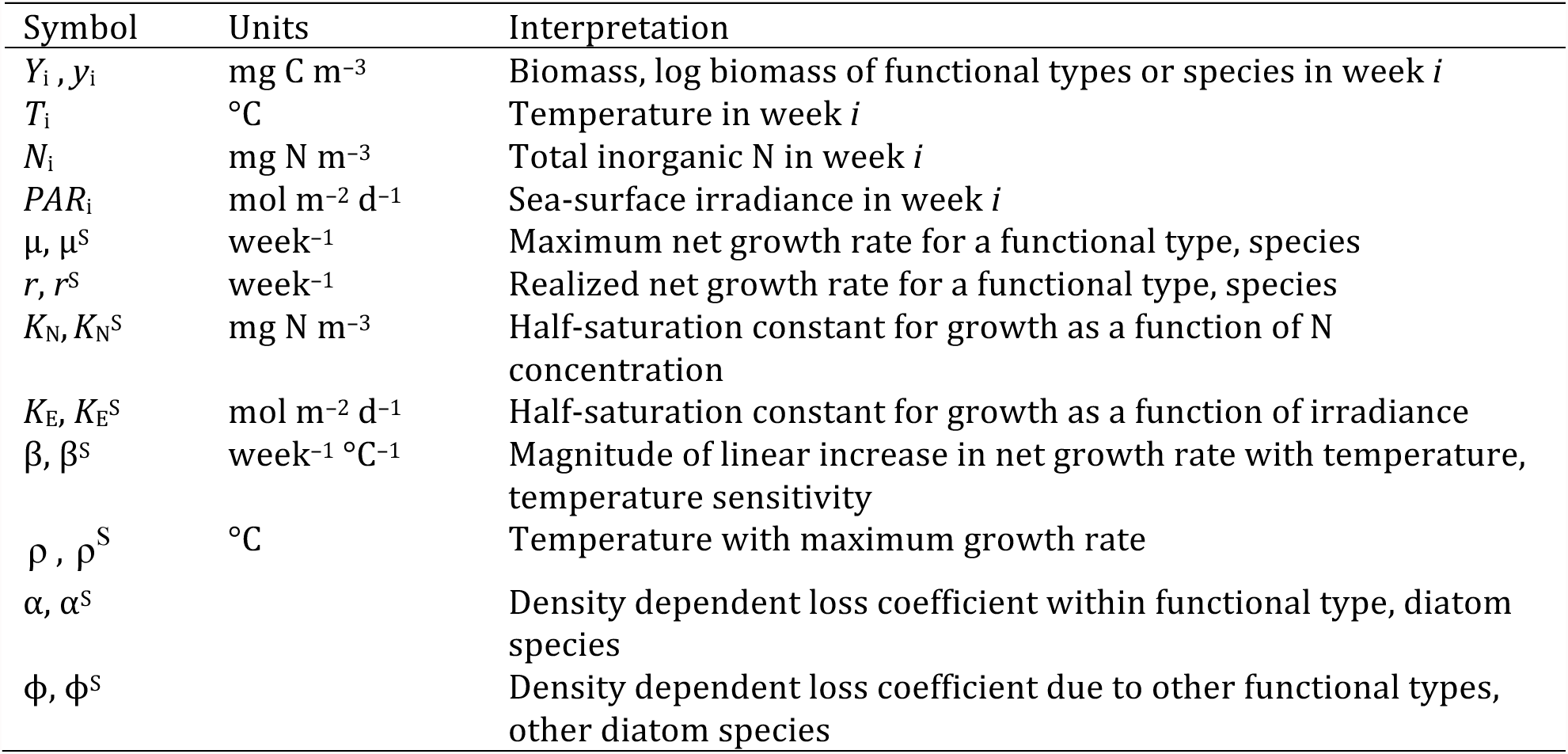
Key symbols for data and traits in the models. Traits for diatom species (as opposed to functional types) have a superscript *S* added.

| Symbol | Units | Interpretation |
| --- | --- | --- |
| $Y_i, y_i$ | mg C m<sup>-3</sup> | Biomass, log biomass of functional types or species in week *i* |
| $T_i$ | °C | Temperature in week *i* |
| $N_i$ | mg N m<sup>-3</sup> | Total inorganic N in week *i* |
| $PAR_i$ | mol m<sup>-2</sup> d<sup>-1</sup> | Sea-surface irradiance in week *i* |
| $\mu, \mu^S$ | week<sup>-1</sup> | Maximum net growth rate for a functional type, species |
| $r, r^S$ | week<sup>-1</sup> | Realized net growth rate for a functional type, species |
| $K_N, K_N^S$ | mg N m<sup>-3</sup> | Half-saturation constant for growth as a function of N concentration |
| $K_E, K_E^S$ | mol m<sup>-2</sup> d<sup>-1</sup> | Half-saturation constant for growth as a function of irradiance |
| $\beta, \beta^S$ | week<sup>-1</sup> °C<sup>-1</sup> | Magnitude of linear increase in net growth rate with temperature, temperature sensitivity |
| $\rho, \rho^S$ | °C | Temperature with maximum growth rate |
| $\alpha, \alpha^S$ | | Density dependent loss coefficient within functional type, diatom species |
| $\phi, \phi^S$ | | Density dependent loss coefficient due to other functional types, other diatom species |

The traits to be estimated appear in the multiplicative growth rate. The density-independent component of the growth rate for functional type *i* from week *w*-1 to week *w, r_i,w_* depends on Michaelis-Menten functions of irradiance (*PAR*, mol m^−2^ d^−1^) and nitrogen concentration (*N*, μmol L^−1^), and a function of temperature (*T*, °C), according to
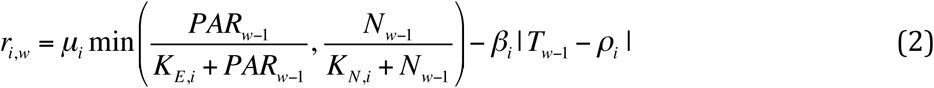

where *ρ_i_* denotes the optimum growth temperature for the biomass of functional type *i* and *β_i_* > 0 is a temperature sensitivity parameter quantifying the increase in the density-independent growth rate *r_i,w_* for a 1°C change in temperature towards the optimum temperature *ρ_i_* and *vice-versa*. Saturating functions of irradiance and nutrient concentration and their combination with a minimum function are commonly used to moderate growth rate (Denman & Peña 1999, Healey et al. 2009). The net growth rate *μ_i_* > 0 is the density-independent growth rate of the *i*^th^ functional type at optimal temperature, irradiance and nitrogen concentration. The effects of irradiance and nitrogen concentration on the growth rate are represented by saturating functions parameterized by the half-saturation constants *K*_E,i_ >0 and *K*_N,i_ >0 representing respectively the irradiance level and nitrogen concentration at which the net growth rate at optimal temperature drops to, *μ_i_* / 2. The Michaelis-Menten saturating functions are combined with a minimum function so that only the most limiting resource affects growth rate at a time, according to Liebig’s law of the minimum (van der Ploeg & Kirkham 1999). During model development, we explored the possibility of a multiplicative interaction between light and nutrients, but we found the results to be more difficult to interpret.

To accommodate density-dependent factors including grazing, viral attack, aggregation and sinking, we introduce density dependent loss terms. In the absence of direct observations of these losses, we parameterize the density-dependent losses with *α_i_* and *ϕ_i_* to quantify the feedbacks on the growth rate of the *i*^th^ functional type from its own biomass and from the combined biomass of the other functional types in the community, respectively. The terms involving *α_i_* and *ϕ_i_* distinguish two different density-dependent loss terms, which could result from specialist and generalist grazer populations, respectively. For purposes of estimating the parameters in the model, we rewrote Eq. (1) on the natural logarithmic scale as
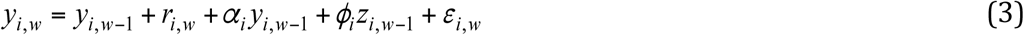

where *y_i,w_* = log(*Y_i,w_*), *z_i,w_* = log(*Z_i,w_*), and *ε_i,w_* = log (*η_i,w_*)

We adapted the functional-type level model described above to define traits at the species level. This task was challenging for two reasons namely, the greatly increased number of parameters to be estimated and the fact that most species are absent from the time series for most of the time, either because they were absent or their abundance was below the detection limit. By contrast, missing values were rare in the time series of functional type biomasses. We restricted the species-level analysis to the 10 diatoms that were observed in about half of the sampling occasions or more. These species may not be representative of the functional type dynamics as a whole because the selected species only represent 11% of the diatom functional type biomass. In order to estimate a growth rate, biomass observations for any particular species must be available on numerous pairs of successive weeks. We extracted pairs of observations from the full time series to estimate the growth rate from week *w-1* to *w*, conditional on the species being observed during weeks *w-1*. The species-level model differed from Eqns. (1-3) only in the definition of the biomass terms *Y_i,w_* and *Z_i,w_* and the interpretation of the density-dependent terms *α* and *ϕ*. To emphasize the differences between the functional type and species-level models, we have added a superscript *S* to the notation for each trait in the species-level model. In the species model, *Y_i,w_* was the biomass of species *i* in week *w*, and *Z_i,w_* was the sum of the biomass of all species in the same functional type as species *i*, except for species *i*, in week *w*. The density-dependent parameter *α* reflects the effect of species *i* on itself while *ϕ* describes the density-dependent effect due to all species in the same functional type as species *i*, except for species *i*.

The model is developed with a Bayesian approach (Gelman et al. 2013). Briefly, Bayesian analysis departs fundamentally from classical statistical methods in that it treats any unknown quantity *θ* as a random variable. As a results, Bayesian inference requires the specification of a probability distribution *p*(*θ*) called prior distribution to describe the uncertainty about plausible values of 9 before taking the data into inconsideration. Upon observing the data, *y*, the likelihood function *p*(*y* | *θ*) of the data is combined with the prior distribution *p*(*θ)* to produce the posterior distribution *p*(*θ | y*) which results from Bayes’ rule as
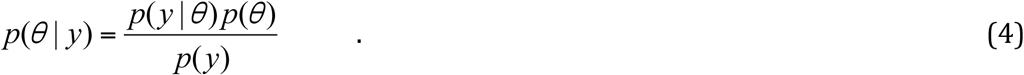

The quantity *p*(*y*) that appears in the dominator of Eq. (4) is the marginal distribution of the data, defined as 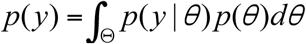 which is nothing but a normalizing constant required to make *p*(*θ* | *y*) integrate to 1 over the parameter space Θ so that it is a proper probability distribution. Therefore Eq. (4) can be written as *p*(*θ* | *y*) ∝ *p*(*y* | *θ*)*p*(*θ*), where *α* stands for “proportional to”.

The posterior distribution *p*(*θ* | *y*) represents the data-updated information and as such, is the basis of Bayesian inference about unknown quantities including model parameters, missing values, and yet unseen data (prediction). Having an entire distribution rather than mere point estimates allows one to fully account for uncertainty. Bayesian conclusions are essentially probability statements based on the posterior distribution. All Bayesian computations are based on probability rules, resulting in more intuitive statements than counterparts in classical statistics.

The main problem of Bayesian inference comes from the difficulty in evaluating integrals like the one in the denominator of Eq. (4). In most practical cases the posterior is not available in closed-form, and sampling-based algorithms, mostly Markov chain Monte Carlo (MCMC) methods (Gilks et al. 1996; Gilks 2005) are typically used to simulate from it and base posterior inferences on the simulated samples. MCMC methods indirectly simulate from a distribution *g* when direct simulation from it is difficult or impossible. The rationale of MCMC sampling is to set up a Markov chain whose stationary distribution is the distribution *g* of interest, in this case the joint posterior distribution *p*(*θ* | *y*).

Consequently, simulation of *θ*^(1)^, *θ*^(2)^,… from the chain yields a series with the property that for large enough *j*, the density of *θ*^(*j*)^ is approximately *g*. In other words, for a large enough “burn-in” period *n*, *θ*^(*n*+1)^, *θ*^(*n*+2)^,… can be regarded as a dependent series with marginal density *g*. Therefore, empirical moments of this series yield approximations of the moments of *g*. In dealing with complex problems, an extension of the model beyond the simple likelihood-prior-posterior scheme is often required, yielding hierarchical Bayesian (HB) models (Gelman et al. 2013).

The model fitting to the functional group biomass data was based on independent priors defined to be fairly uninformative for most parameters. More specifically, we assigned standard normal priors to the density-dependence parameters *α_i_* and *ϕ_i_*, normal priors centered at 13°C (the average temperature at L4 Station over the time series) with variance 10 to the optimal temperatures *ρ_i_*, standard normal priors constrained to positive values on the temperature sensitivities *β_i_*, Gamma priors with mean 2 and variance 1 to the functional group-specific net growth rates, *μ*, Gamma(1,1) priors on the functional group specific error variances, and positively truncated normal distributions centred at zero with variances 100 and 0.1 on the irradiance and nitrogen half-saturation constants *K*_E_ and *K*_N_, respectively. We imposed relatively informative priors on *μ* and *K*_N_ based on our previous experience with the L4 dataset to facilitate model identifiability. For the species model, we defined the prior distributions on *μ*, *K*_E_, *K*_N_, and *β* to be concentrated around the FG estimates with relatively small variances.

Since the joint posterior is not available in closed-form, we used Markov-chain Monte-Carlo (MCMC) methods (Gilks et al. 1996) implemented in OpenBUGS (Thomas et al. 2006) to simulate from it. We ran 40 000 iterations of two parallel Markov chains starting from dispersed initial values, discarded the first 15 000 samples from each Markov chain as burn-in period and thinned the remaining 25 000 samples by factor of 25. We assessed the convergence of the Markov chains through visual inspection of traceplots and autocorrelation functions. For all parameters, the Markov chains mixed well with the sampler jumping freely around the parameter space as illustrated by Figs. S1-S3 in the Supplementary Material. We also conducted a simulation study to evaluate performance of our model in terms of inference and prediction. We extracted from the L4 functional group biomass data the biomass of the two functional groups with complete data over the time series namely, diatoms and phytoflagellates. We fitted our model to the data with all environmental variables (temperature, irradiance and nitrogen concentration) set to the observed values at Station L4 over the time series. We considered the posterior predictive means as simulated data from the hypothetical two functional group system under our model, with underlying parameter values given by the posterior mean estimates. We fitted the model back to the simulated data. The model was effective at retrieving the underlying parameter values as indicated by Figs. 4S and 5S in the Supplementary Material.

## Results

The three environmental drivers of phytoplankton growth rate included in this study (temperature, irradiance, and nitrogen concentration) exhibit strong, regular seasonal oscillations over the seven-year time series (Widdicombe et al. 2010, Mutshinda et al. 2016). The phytoplankton biomass for each of four functional types each exhibit distinctive patterns of intra-annual variation (Fig. 1). Diatoms bloom first, increasing steadily in biomass from day 60 to day 180. Dinoflagellates and coccolithophorids bloom slightly later, reaching a maximum biomass at approximately day 225. The amplitude of dinoflagellate biomass is the greatest across the four types and their sustained maximum growth and loss rates are also the largest. Phytoflagellates have the least inter-annual variability, with two minor biomass peaks at approximately day 110 and day 215. Our model was able to reproduce the temporal patterns in the biomass of all functional groups with narrow posterior predictive intervals relative to the total variation in the data (Fig 1). There was insufficient temporal resolution in the data to observe short-term acclimation to changing conditions, so our focus remained on steady-state traits similar to those usually used in phytoplankton community models.

**Figure 1.**
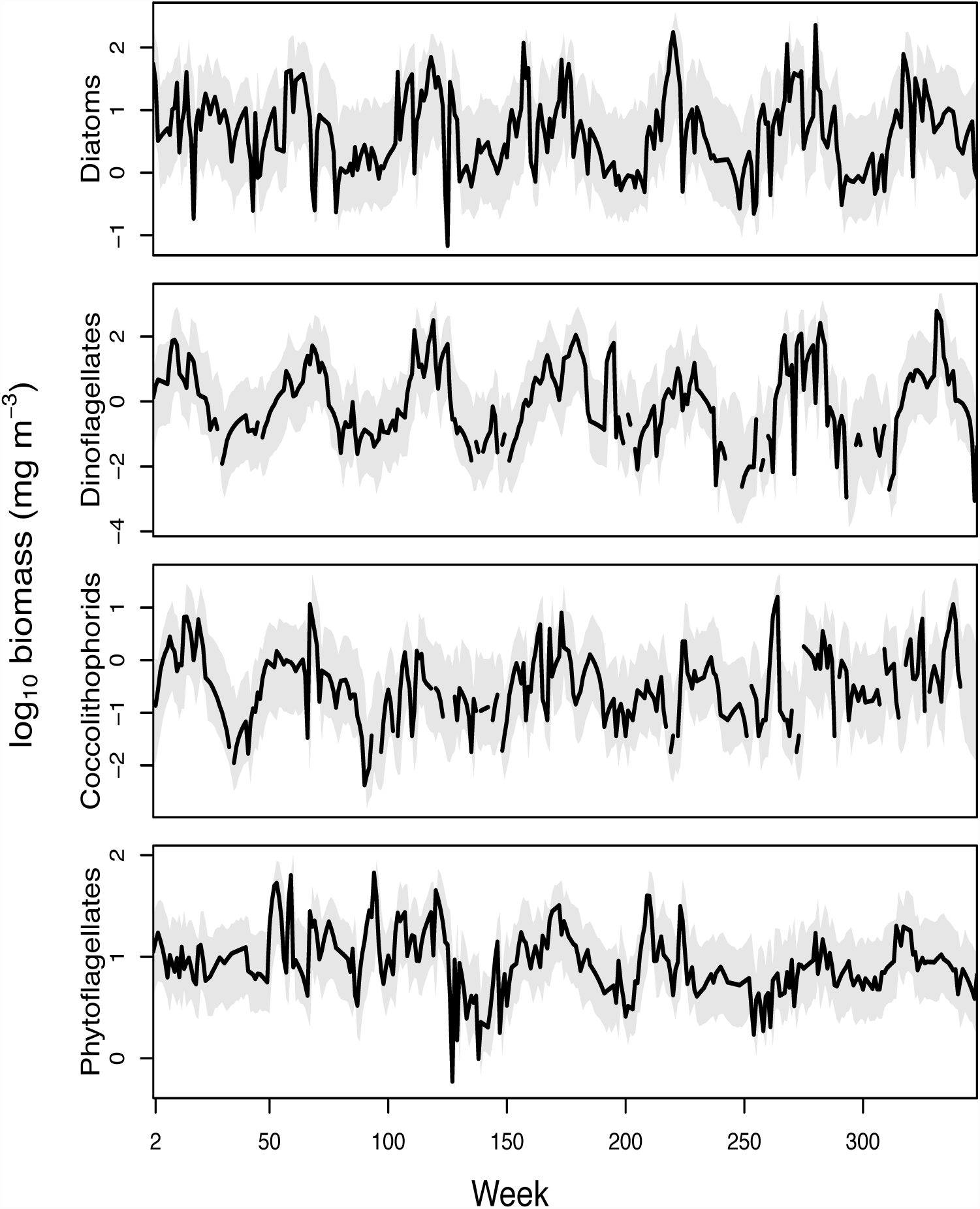
Observed (black line) and predicted (shaded region) log10 carbon biomass (mg C m^−3^) of each functional type (diatoms, dinoflagellates, coccolithophorids, and phytoflagellates) at Station L4. The prediction region is the 95% credible range of biomass from the functional type model.

### Functional-type level analysis

The maximum net growth rate trait, *μ_i_*, is the largest growth rate of functional type *i* under any irradiance and nutrient conditions, at its optimal temperature for growth, not including density-dependent grazing, but incorporating linear grazing rates. There is substantial variability in maximum net growth rates between functional types (whiskers on Fig. 2a). As a group, diatoms have the largest net growth rate with mean doubling time 2.5 days, followed by dinoflagellates and coccolithophorids with mean doubling times 3.5 and 4 days, respectively. Phytoflagellates have the lowest net growth rate with approximate mean doubling time 5 days.

**Figure 2.**
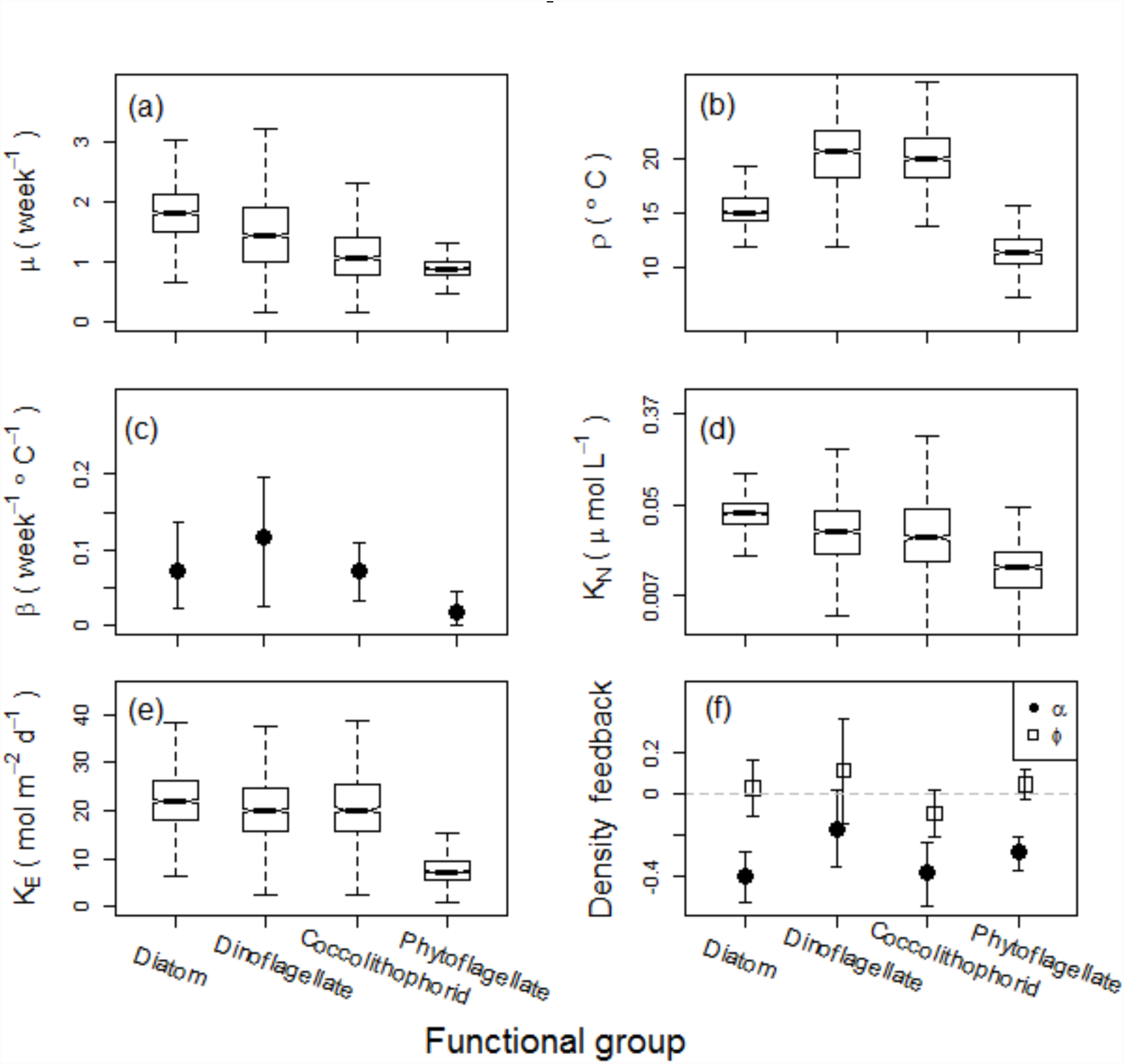
Posterior distributions of traits governing growth rate at the functional type level. (a) Maximum net growth rate, *μ* (week^−1^), (b) the optimal temperature for growth ρ (°C), (c) temperature sensitivity, β (week^−1^ °C^−1^), (d) the half-saturation constants for nitrogen, *K*_N_ (μmol L^−1^), (e) the half-saturation constants for irradiance, *K*_E_ (mol m^−2^ d^−1^), and (f) density-dependent effects on the growth rate of each functional type attributed to their own biomass (*α*, solid circle) and to the total biomass of the other functional types in the community (*ϕ*, open squares). Box plots show median (thick line), the interquartile range (box) and the full range of the data or 1.5 times the interquartile range, whichever is smaller (whiskers). In panels (c) and (f) error bars indicate 95% credible intervals on the posterior means and are used because posterior distributions are approximately normal. In (f), the horizontal dashed line indicates no density-dependence. The vertical scale in (d) is logarithmic to facilitate the display of the wide range of values.

The estimated optimal temperatures for growth for diatoms, dinoflagellates, coccolithophorids and phytoflagellates are 15°C, 20°C, 20°C and 11°C, respectively, implying that higher temperature conditions would favor dinoflagellates and coccolithophorids biomass accumulation (Fig. 2b). As a group, dinoflagellates are the most responsive to temperature changes with a temperature sensitivity parameter roughly twice those of diatoms and coccolithophorids. On the other hand, phytoflagellates are essentially insensitive to temperature changes at Station L4 with temperature sensitivity parameter estimated to be close to zero (Fig. 2c).

The nitrogen (nitrate, nitrite, plus ammonia) half-saturation constants, *K_N_*, for all groups are comparable to those found in lab studies and used in models (Fig. 2d). Phytoflagellates have the smallest half-saturation constants for irradiance, which is consistent with their relatively small amplitude of biomass variation over the time series. The irradiance half-saturation constants for the other three functional groups are not credibly different from one another (Fig. 2e).

The half-saturation constants for nitrogen concentration (posterior means ranging from 0.008 to 0.04 μmol L^−1^) are quite close to the minimum values of the corresponding environmental data observed over the time-series (0-15 μmol L^−1^), suggesting that this trait may not be informative for predicting the biomass growth rate of these functional types at this location for most of the year. Conversely, the half-saturating constants for sea-surface irradiance (posterior medians ranging from 8 to 23 mol m^−2^ d^−1^) span most of the lower half of the inter-annual variation in irradiance (10-50 mol m^−2^ d^−1^), indicating that phytoplankton growth rates vary with irradiance (light is sub-saturating) for much of the year (Fig. 2e).

All four phytoplankton functional types are affected by density-dependent loss rates (Fig. 2f). These losses have the largest effect at high biomass concentrations and can explain the maximum biomass concentration for each functional type, but they are also active at low biomass concentrations and are responsible for decreases in biomass when growth conditions are unfavorable. Density-dependent losses are a combination of grazing, viral attack, and aggregation and sinking following bloom collapse. For each functional type, we distinguished between density-dependent feedback due to the functional type’s own biomass (*α*) and the feedback due to the aggregate biomass of all the other functional types (*ϕ*). If the density-dependent loss terms are primarily due to grazing, we could interpret *α* as representing the losses due to grazers specializing on one functional type and *ϕ* as representing losses due to generalist grazers supported by populations of the other functional types. Since *α* < 0 for all functional types (and *ϕ* ≈ 0), the biomass of each functional type is largely regulated by specialist grazers and generalist grazers have weak density-dependent effects.

### Species-level analysis

Diatom species’ net growth rates were smaller than the functional type counterpart for all ten species examined (Fig. 3a). For all species, the half-saturation constants for nitrogen were roughly twice as large as the functional group estimates, and the temperature sensitivity parameters (the *β*^S^) were close to 0.10 week^−1^ °C^−1^ (which is within the 95% credible interval of functional group estimate), except the two *Pseudo-nitzschia* strains which stood out with temperature sensitivity parameters twice as large (Fig. 3b). The optimal growth temperatures were extrapolated outside the range (7.5-18.8°C) of observed temperatures for most species (not shown). The species’ half-saturation constants for irradiance (not shown) varied from 25 to 30 mol m^−2^ d^−1^. For the species model, the density dependent loss terms were redesigned to identify species-specific density-dependent loss rates and generic functional type density-dependent loss rates. The posterior distributions of density dependent parameters *α^S^* and *ϕ^S^* (Fig. 3d), imply a stronger negative feedback on each diatom’s biomass growth from its own biomass than that from the combined biomass of other diatoms *(a^s^* < *ϕ^s^*), consistent with niche differentiation within functional types (Mutshinda & O’Hara 2011). Some of the *α^S^* and *ϕ^S^* for diatoms were positive, suggesting the presence of mutually beneficial or commensal effects in some species.

**Figure 3.**
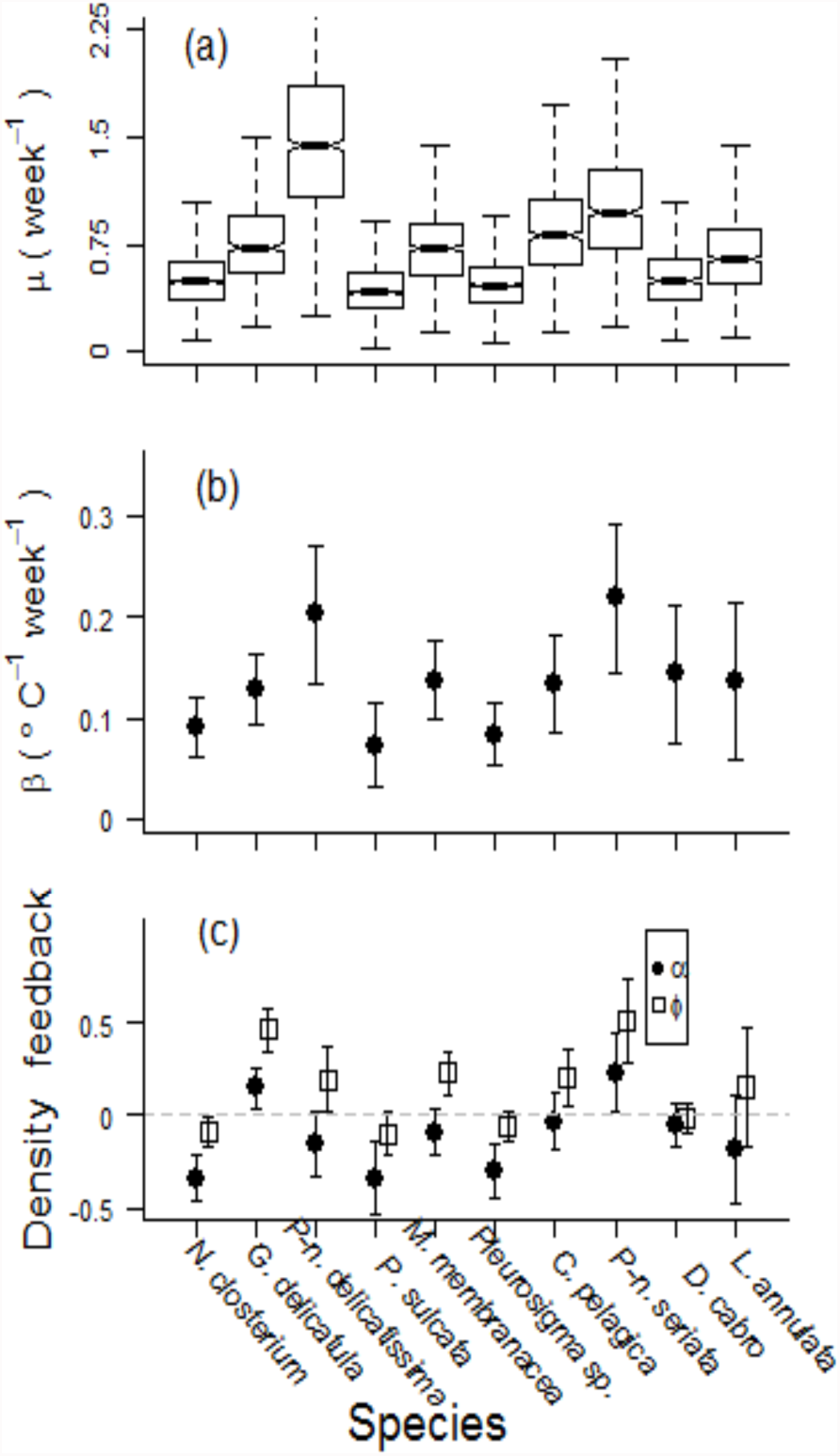
Species-level trait values for ten diatom species. Species are arranged in order of the number of weeks they are present in the time-series from *Nitzschia closterium* (93% of weeks) to *Lauderia annulata* (48%). (a) Maximum net growth rate, *μ^S^* (week^−1^), (b) temperature sensitivity, β (week^−1^ °C^−1^) (c) density-dependent effects on the growth rate of each species attributed to their own biomass (*α^S^*, solid circle) and to the total biomass of the other species in the same functional type (*ϕ^S^*, open squares). Box plots show median (thick line), the interquartile range (box) and the full range of the data or 1.5 times the interquartile range, whichever is smaller (whiskers). In panels (b) and (c) error bars indicate 95% credible intervals on the posterior means and are used because posterior distributions are approximately normal.

## Discussion

Trait-based models of phytoplankton productivity promise to deliver robust projections of phytoplankton community dynamics under future climate scenarios. Phytoplankton traits are estimated in the lab one species at a time but are commonly aggregated into functional types for ocean biogeochemical models (Anderson 2005, Le Quéré et al. 2005, Litchman et al. 2006). There are several challenges that arise in the estimates of phytoplankton traits for trait-based models. Most species in diverse communities have not been systematically studied in the lab. Trait values vary across species, even within functional types, and it is not clear how to produce an average trait value for modeling functional types. In addition, there is considerable phenotypic plasticity in traits. Furthermore, grazing rates, viral and parasitic loss rates, sinking rates and biotic interactions, such as allelopathy or mutualisms, can be complex and highly variable from species to species. It is difficult to get good estimates of loss terms, such as grazing rate and viral lysis, that are inherently species specific and patchy in time and space, and we are just starting to learn about the consequences of the many, complex biotic interactions between phytoplankton and their microbial communities (Sher et al. 2011, Amin et al. 2015). It may be possible to overcome some of these myriad challenges using phytoplankton traits estimated directly from field data or by combining lab-based traits with niches estimated from the field (Edwards 2016). Here we extract functional type and species-level phytoplankton traits from time-series data from a well-studied coastal temperate phytoplankton community in the Western English Channel (Harris 2010, Widdicombe et al. 2010). The variability in trait values we extracted from field data likely reflects in part true variability due to acclimation of species within communities to changing environmental conditions and changing community composition through the seasons. While some of the traits estimated here are consistent with laboratory estimates based on single species analyses, many are not, indicating more work is needed to understand how phytoplankton respond in natural communities.

Our estimates (posterior means) of maximum net growth rate for the phytoplankton functional types range from 0.9 to 1.8 week^−1^ (a mean doubling time of 5 to 2.5 days) and for the 10 individual diatom species from 0.5 to 1.5 week^−1^ (a mean doubling time of 10 to 3 days). Our growth rate estimates are consistently lower than lab-based estimates of growth rate from unialgal cultures and *in situ* field estimates of growth rate of individual species (grazers excluded) that can double more than once a day (Furnas 1990, 1991, Raven et al. 2005). Maximum *in situ* growth rates for three of our ten diatom species have been estimated from daily counts during April in the Irish Sea: *Pseudo-nitzchia* sp., 0.24 d^−1^; *Guinardia delicatula*, 0.18 d^−1^; *Lauderia annulata* 1.42 d^−1^ (McKinney et al. 1997). Their growth rate estimates are significantly larger than ours. Weekly counts, used in our study, are likely to lead to smaller maximum net growth rates than daily counts because the coupling between growth and loss processes will be tighter when averaged over a week instead of a day. Additionally, we expect our estimates of maximum growth rates to be lower than traditional estimates of individual species growth rates in the lab and field because our growth rate estimates include linear loss terms due to grazing, viral and parasitic loss and are therefore similar to a net phytoplankton community growth rate. Our values for net growth rate are consistent with satellite-based estimates of monthly median phytoplankton growth rates in temperate regions with strong seasonal blooms, 0.35 to 4.2 week^−1^ (Westberry et al. 2008). Microzooplankton grazing at Station L4 and elsewhere has been estimated to account for about two-thirds of phytoplankton growth (Fileman et al. 2002, Calbet & Landry 2004, Chen et al. 2009, Bernard et al. 2012). Given our estimates of maximum growth rates tend to be much lower than estimates of growth rate from lab studies, this suggests loss rates due to grazing and parasitoid and viral attack may be higher than often assumed.

It would be plausible for there to be no relationship between our field based estimates of maximum growth rates across the functional types even if there are differences in maximum net growth rate since the grazing and other linear loss terms represent such a large fraction of maximum net growth rate. We find the rank order in our estimates of net growth rates for the functional types (diatoms > dinoflagellates > coccolithophorids >phytoflagellates) are generally consistent with growth rates reported from laboratory culture work and field observations (Furnas 1991, Cermeño et al. 2005, Raven et al. 2005, Laws 2013). In the Western English Channel, we find diatoms have the largest maximum net growth rate followed by dinoflagellates and coccolithophorids, whereas phytoflagellates have the smallest net growth rate (Fig. 2a). These results indicate that lab-based maximum growth rates combined with a constant loss rate used by many models may be a reasonable proxy for net growth rates in natural communities.

The effect of temperature on phytoplankton species growth rates is commonly described using the Q_10_ approximation, which is the multiplicative effect of a 10°C change in temperature on growth rate. This value is typically about 2, ranging from 1.88 to 2.3 for phytoplankton (Eppley 1972, Bissinger et al. 2008). The range of temperatures at Station L4 (about 8-19°C) is narrow compared to the width of many phytoplankton temperature niches (Irwin et al. 2012, Boyd et al. 2013), so we used a linear model to describe the effect of temperature on growth rate (see Montagnes et al. 2003 for additional rationale for using a linear model). The temperature sensitivity of the functional types, β, is about 0.12 week^−1^ °C^−1^ for dinoflagellates, 0.07 week^−1^ °C^−1^ for diatoms and coccolithophorids, and 0.02 week^−1^ °C^−1^ for phytoflagellates (Fig. 2c). These estimates suggest that on average, growth rate would increase from 1 week^−1^ to roughly 1.72 week^−1^ for diatoms,1.42 week^−1^ for diatoms and coccolithophorids and 1.12 week^−1^ for phytoflagellates, with an increase in temperature of 6°C (half the annual amplitude in temperature), starting and ending below their temperature optima. Analysis of the change in maximum growth rate with temperature from unialgal lab cultures (Montagnes et al. 2003) found slopes of 0.11 to 0.54 week^−1^ °C^−1^ for dinoflagellates, consistent with the posterior mean (0.12 week^−1^ °C^–1^) found in this study (Fig. 2c), and 0.084 to 0.97 week^−1^ °C^−1^ for diatoms, which is larger than the posterior mean (0.07 week^−1^ °C^−1^) found here. For phytoflagellates, our estimate of the temperature sensitivity trait, β, is approximately 0.02 week^−1^ °C^−1^ which is close to zero with a narrow credible interval, so we conclude that temperature has essentially no effect on the growth rate of this functional type at this site. Possible interpretations for this result are that the phytoflagellates have broad temperature optima for growth rate or the functional type is composed of many species with specialized optimal growth temperatures spread across the range of observed temperatures (Eppley 1972, Boyd et al. 2013). This does not appear to be the case for the other three functional groups, even if there is species turnover during the year, there is still a fairly strong imprint of temperature on the growth rate of the functional type as a whole. An alternative explanation is that an increase in water column stability favors dinoflagellate and coccolithophorid biomass accumulation (Margalef 1978, 1997). The optimal temperature for growth at the functional type level varies as expected. Phytoflagellates have the lowest and coccolithophorids and dinoflagellates the highest optimal temperatures for growth. However, the optimal temperature for dinoflagellates which, as coccolithophorids bloom later than diatoms in the season, exhibit more variability (Fig. 2b) implying that dinoflagellates have a wider temperature niche than the other functional groups. Since temperature is correlated with stability and we don’t have an independent measure of stability, our model is unable to distinguish between the direct effects of temperature and the effect of water column stability on the growth rate of phytoplankton.

Temperature optima estimates for individual diatom species were not identified within the range of observed temperatures. We interpret this result as consistent with wide temperature response curves, relative to the narrow temperature range at Station L4, for the species under study (Boyd et al. 2013). The estimated temperature sensitivity parameters for individual diatom species (β^S^) are larger than the functional type counterpart (Fig. 3b), which is to be expected as aggregating species into functional types should reduce the effective strength of temperature on growth rate averaged over the portion of the community belonging to each functional type.

Light and nitrogen limitation of net growth rate is determined by Michaelis-Menten half-saturation trait values, *K*_N_ and *K*_E_. Three sources of inorganic nitrogen: nitrate, nitrite, and ammonium are considered in our estimate of *K*_N_ for inorganic nitrogen. As a result our estimate of *K*_N_ at the functional type level is largely determined by the inorganic nitrogen. Nitrogen half-saturation constants for phytoplankton species can vary from 0.08 to 8.4 μmol L^−1^ in the lab (Litchman et al. 2006). Our estimates (posterior means) of nitrogen half-saturation constants for individual diatom species, many with large cell size, ranged from 0.08 to 0.12 μmol L^−1^ which is with this range. Our values for functional types range from about 0.008 to 0.04 μmol L^−1^ and are either on the lower end or smaller than typical literature values for unialgal cultures. The half saturation constants for inorganic nitrogen for functional types at this site are also low relative to all nitrogen concentrations observed in seawater at this site (ranging from 0.1-16 μmol L^−1^), indicating that nitrogen limitation is only a significant factor affecting growth rates of functional types, particularly diatoms and dinoflagellates, in the warmest part of the summer. One reason *K*_N_ may be lower in the field relative to laboratory studies is that organic nitrogen may be an important source of nitrogen for some species, particularly the dinoflagellates and phytoflagellates, but also some diatoms such as *Pseudo-nitzschia delicatissima* (Loureiro et al. 2009). If organic sources are important for these groups, for example following the crash of a diatom bloom when inorganic nitrogen concentrations are low, estimated *K*_N_ may be artificially low since organic sources were not included in the model. Alternatively, since nitrogen is taken up rapidly when available, bulk estimates of reactive nitrogen concentration sampled weekly may be relatively uninformative at physiological scales (Laws 2013). The phytoflagellates have the lowest *K_N_* of approximately 0.008 μmolL^−1^, which is much less than the values for the other three functional groups, namely 0.03 μmolL^−1^ for dinoflagellates and 0.04 μmolL^−1^ for diatoms and coccolithophores. The phytoflagellates category is taxonomically diverse, but over half the biomass is found in unidentified cells smaller than 5μm in diameter. The most significant feature of our results is that the phytoplankton dynamics at Station L4 is consistent with very low *K*_N_ compared to values estimated from laboratory cultures (Litchman et al. 2007). The *K*_N_ at Station L4 are 5-10 fold smaller than half-saturation constants for nitrate often employed in ecosystem models (Gregg et al. 2003, Merico et al. 2004). The intermediate complexity marine ecosystem model constructed by Moore et al (2002) is an exception; this model uses a very low *K*_N_ for ammonium of 0.004 μmolL^−1^ for small cells, much lower than our values for Station L4 (Moore et al 2002). Generally *K_N_* values for ammonium are smaller than for nitrate (Litchman et al. 2007; Merico et al. 2004).

Light limitation is frequently parameterized by a half-saturation coefficient, *K*_E_, or the irradiance at which light saturates growth, *E*_k_. For comparison between the two, we divide E_k_ by 2 to roughly approximate *K*_E_. In natural populations in coastal regions, *E*_k_ varies from 40-500 μmolm^−2^ s^−1^ (Kirk 2010), corresponding to *K*_E_ of about 2-22 mol m^−2^d^−1^. Estimates of *K*_E_ in unialgal cultures range from 3.5-7.8 mol m^−2^ d^−1^(Litchman et al. 2006), and varies with steady state irradiance (Gregg et al. 2003, Kirk 2010). At Station L4, our estimates of *K*_E_ for functional groups range from 8 to 23 mol m^−2^ d^−1^, but these are based on sea-surface irradiance and thus are larger than they would be based on average *in situ* irradiances. Individual diatom species have *K*_E_ ranging from 25 to 30 molm^−2^ d^−1^. These results suggest that irradiance at Station L4 is limiting for diatoms, dinoflagellates and coccolithophorids during much of the year, since sea-surface PAR ranges from 10-50 molm_−_2 d^1^ and only exceeds *E*_k_ ≅ 2*K*_E_ ≅ 40 mol m^−2^ d^−1^ for these groups during short periods in the summer. By contrast, phytoflagellates have *K*_E_ near the minimum levels of PAR and so they experience saturating irradiance for most of the year. One possible hypothesis is that their small size confers a low pigment package effect, meaning they have high light absorption per unit of pigment, giving them an advantage over functional types with larger cells under low light conditions (Finkel & Irwin 2000, Finkel 2001, Finkel et al. 2004). Furthermore, if some of the phytoflagellates use alternative energy sources, they may require less chlorophyll and be less sensitive to changes in irradiance. While some dinoflagellates are known to be heterotrophic and mixotrophic (Stoecker 1999), unlike phytoflagellates their growth rate is strongly affected by low temperatures in winter, reducing their growth rate in winter relative to phytoflagellates (Fig. 1). Phytoflagellates appear to be able to acclimate to very low light, giving them a competitive advantage over other functional types, especially in winter.

Many studies of zooplankton grazing focus on the linear grazing rate (Landry & Hassett 1982, Calbet & Landry 2004, Zheng et al. 2015), which in our model is combined with gross phytoplankton growth rate to obtain the maximum net growth rate trait, *μ*, which is assumed to be constant for each phytoplankton functional type. More complex formulations of zooplankton grazing rates permit diel and seasonal variation in grazing rates and non-linear grazing rates (Tsai et al. 2005) or describe prey switching or selectivity by grazers (Gentleman et al. 2003, Vallina et al. 2014), but we do not consider these mechanisms. Our model incorporates density-dependent loss terms to describe consumption of phytoplankton by grazers along with other loss processes.

All four functional types exhibit strong density-dependent loss. Assuming the loss term is primarily attributable to grazing, all functional groups are primarily grazed by specialists (*α* < 0, Fig. 2). However, while diatoms, dinoflagellates and phytoflagellates are virtually unaffected by generalist grazers (*ϕ* ≅ 0), coccolithophorids are affected by both specialist and generalist grazers. Each functional group is more negatively affected by its own biomass than by the combined biomass of other functional groups, which provides evidence for niche differentiation between functional groups.

The results at the species level are more variable. Five of our ten diatom species (*Guinardia delicatula*, *Pseudo-nitzschia delicatissima*, *M menbracea*, *C*. *Pelagica* and *Pseudonitzschia seriata*) exhibit positive density-dependent effects (*ϕ^S^* > 0, Fig. 3c) with increased biomass of all other diatoms. This could be an indication that these species experience less grazing pressure when the biomass of other diatoms is high (“kill the winner”, Vallina et al 2014). Two species, *Guinardia delicatula* and *Pseudo-nitzschia seriata*, have positive density-dependent effects resulting from their own biomass (*α^S^* > 0), indicating that increases in their biomass can increase their own growth rates. Many strains of *Pseudonitzschia* have been shown to produce the neurotoxin domoic acid (Bates et al. 1998, Fehling et al. 2004), suggesting this positive density-dependence may be a result of allelopathy, although *G*. *delicatula* does not produce toxins and *Pseudo-nitzschia delicatissima* has *α^S^* < 0. Finally, three species *Nitzschia closterium, P. sulcata* and *Pleurosigma* sp. have strong negative density-dependent feedbacks from their own biomass (*α^S^* < 0). *Nitzschia closterium* is known to produce mucus that may increase its export at high densities, which is consistent with this result (Najdek et al. 2005).

Our analysis of ten diatom species demonstrates the potential and challenges of this approach for determining trait values and modeling dynamics of individual species. These species were the most frequently observed in the population, but account for only 11% of the total biomass, on average. Species with fewer observations are less likely to yield informative estimates of trait values due to a lack of data, but account for the vast majority of the biomass. Since our ten species sample is a minority component of the diatom community and represents species present much of the year in contrast to species present for only a few weeks at a time, there is no reason to expect the trait values of these species to be representative of the functional type as a whole. In fact, we observed systematic differences between trait values for these species and the diatom functional type: the species-specific maximum growth rates are lower and the half-saturation constants for light and nitrogen are higher relative to the functional group type estimates. Even if we had a random sample of species with trait values representative of the full distribution, determining functional-type level trait values by averaging over species with different traits and changing contributions to the total population can lead to errors due to Simpson’s paradox (Chuang et al. 2009, Williams & Hastings 2011). The uncertainties across the diatom species are large enough to suggest that the trait values may be largely indistinguishable across many species, in particular the irradiance half-saturation constants. An independent analysis showed that diatoms species at Station L4 exhibit neutral dynamics within the diatom functional type most of the time, indicating that predicting biomass dynamics of individual species may be much harder than predicting the dynamics of the aggregated biomass of a functional type (Mutshinda et al. 2016). While it is appealing to estimate trait values for functional types from knowledge of individual species, it may be more prudent to deemphasize species-level details and use realized traits estimated from biomass dynamics aggregated to the functional-type level.

## Conclusions

This study enables a comparative analysis of trait values used in biogeochemical models of phytoplankton communities and the trait values estimated from lab studies on individual phytoplankton. The realized traits we quantified could be different from those estimated in the lab because they are functional-type level aggregates and include factors such as phenotypic plasticity and biotic interactions that may vary across species and communities. At Station L4 in the Western English Channel, we found that diatoms have the highest maximum net growth rates, intermediate temperature sensitivity, and high specialist density-dependent loss rates. Dinoflagellates have intermediate maximum net growth rates and high temperature optimum and sensitivity. Coccolithophorids have high temperature optimum, intermediate temperature sensitivity, and are negatively affected by both specialist and generalist density-dependent feedbacks. The phytoflagellates have the lowest maximum net growth rate, low optimum temperatures and sensitivities, and low half-saturation constants for light and nitrogen concentration. The relative differences in maximum net growth rate, specifically the relatively high rates for diatoms, are consistent with differences estimated in the lab and the field, but the absolute magnitudes of the rates are considerably lower because our maximum growth rates include linear loss terms. A comparison of our results with traits estimated in the lab and used in models yields a few insights. Grazing and other linear loss rates, as reflected in a reduction of the gross growth rate, appear be even more important than usually appreciated. We see evidence of complex biotic interactions that are difficult to assess in the lab: all functional types are more susceptible to specialist loss rates, perhaps indicating specialist grazers or viruses. At the species level, there appears to be evidence of species interactions increasing the net growth rate of individual diatom species. The half-saturation constants for nitrogen are lower than typical lab estimates, consistent with the use of a wide range of reactive nitrogen sources and widespread mixotrophy. There is considerable variation in our estimates of the trait values within phytoplankton functional types, which could be due to real physiological changes arising from acclimation to environmental conditions over time, variation across species within a functional type, or a consequence of insufficient data. Time-series of field data combined with our analysis gives us insight into the mechanisms affecting the dynamics of species and whole functional types in natural populations that may improve our ability to scale-up results from species-level studies in the lab to community dynamics in the ocean.

## Acknowledgements

Station L4 phytoplankton biomass and environmental data were provided by the Western Channel Observatory which was funded as part of the UK Natural Environmental Research Council’s National Capability. EMS Woodward and C Harris provided the nutrient data. We thank A Atkinson and T Smyth for assistance and advice. ZVF was supported by the National Science and Engineering Research Council (NSERC) of Canada and the Canada Research Chairs program. AJI was supported by NSERC.

